# Differentiation of KISS1-Expressing Cells from Human Pluripotent Stem Cells: Many Roads To Rome

**DOI:** 10.64898/2026.09.02.748796

**Authors:** Celia Gomez-Sanchez, Nazli Eskici, Shrinidhi Madhusudan, Karoliina Tanner, Kirsi Vaaralahti, Kristiina Pulli, Taneli Raivio

**Author notes:** Corresponding Author: Celia Gomez-Sanchez.

## Abstract

**Introduction:** Kisspeptin-secreting (Kiss) neurons govern human puberty and reproduction. In the arcuate nucleus (ARC), they control pulsatile release of gonadotropin-releasing hormone (GnRH), while Kiss neurons in the preoptic area (POA) control GnRH surge. Since animal models do not fully recapitulate the human phenotype, a human model to study these neurons is crucial.

**Methods:** We differentiated human pluripotent stem cells (hPSCs) into neuron cultures using two distinct strategies: FGF8 protocol, consisting of dual SMAD inhibition (dSMADi), FGF8b, and Notch inhibition; and SHH protocol, consisting of dSMADi and SHH activation, followed by Notch inhibition. Neuron cultures obtained on day 45 were characterized at the mRNA level using RT-qPCR.

**Results:** Both strategies resulted in neuron cultures where significant *KISS1* expression could be detected. SHH-derived neuron cultures expressed high *NKX2-1* and the ARC markers *POMC, NHLH2*, and *NR5A2*, while FGF8-derived neuron cultures expressed low *NKX2-1* and the anterior POA marker *FOXG1*.

**Conclusions:** We provide the first ever strategies to differentiate hPSCs into neuron cultures that express *KISS1*. Future studies providing in depth transcriptomic, protein, and functional characterization are needed to establish the properties of these models.

## Introduction

Human puberty and reproduction are governed by a complex neuroendocrine system orchestrated by hypothalamic kisspeptin-secreting (Kiss) neurons[1]. Kisspeptin (encoded by the *KISS1* gene) activates gonadotropin-releasing hormone (GnRH) neurons, which trigger the release of gonadotropins from the pituitary gland enabling sex hormone production[2]. Two different populations of hypothalamic Kiss neurons can be found, one in the arcuate nucleus (ARC) and one in the preoptic area (POA), also termed anteroventral periventricular nucleus or AVPV in rodents. In the ARC, Kiss neurons form the functional “GnRH pulse generator” unit[3,4]. Kiss^ARC^ frequently coexpress neurokinin B (NKB) and dynorphin (Dyn) in rodents and sheep, and are therefore termed “KNDy neurons”; however, human Kiss^ARC^ neurons more frequently express kisspeptin alone or together with NKB[5]. Meanwhile, Kiss^POA^ neurons control the LH surge[6].

Currently, investigating disorders of human puberty and fertility is challenging due to the lack of human-specific models. While numerous studies have been performed in animals, these often do not fully recapitulate the human phenotype, likely due to differences in estrous cycle and reproduction[5,7,8]. Since direct neuron collection from patients is not possible, a human in vitro model of Kiss neurons is crucial for research in this field.

The ontogeny of Kiss neurons has not been fully elucidated. It has been proposed that a fraction of mouse Kiss^ARC^ neurons may descend from POMC-expressing neural progenitors[9,10]. However, a recent comprehensive work in human pluripotent stem cells (hPSCs)[11] demonstrated that hypothalamic ARC cells differentiated using SHH and WNT activation did not express *KISS1* despite presenting robust expression of *POMC, TBX3*, and *NHLH2*, transcription factors implicated in rodent kisspeptin neuron ontogeny[10,12,13]. Moreover, Kiss^ARC^ and Kiss^AVPV^ differ in their gene expression profile, and it has been suggested that they could have different developmental origins[14]. Indeed, Kiss^AVPV^ could emerge from more anterior neural progenitor cells than Kiss^ARC^, as indicated by the expression of several anterior markers including *Fgfr1, Foxg1, Dlx1*, and *Dlx2*[14], all of which have also been implicated in GnRH neuron development[15–17].

Herein, we report preliminary results from our work showing two differing strategies that enable the differentiation of *KISS1*-expressing neuron cultures from hPSCs, that may display ARC-like and POA-like gene expression signatures.

## Methods

### hPSC culture

The human embryonic stem cell line H9 (WA09 WiCell) was used in this study. hPSC monolayer cultures were plated on Matrigel-coated (Corning, Cat. 356231) 35 mm dishes (ThermoFisher Scientific, Cat. 153066) maintained in mTeSR1 culture media (STEMCELL Technologies, Cat. 85850), which was refreshed daily. hPSCs were passaged after reaching >80% confluency by washing them with PBS (Corning, Cat. 21-040-CV) and dissociating them with 0.5 mM Ultrapure EDTA (Invitrogen, Cat. 15575-038) diluted in PBS. hPSCs were kept at 37°C and 5% CO_2_.

### Kiss neuron differentiation

Neuron differentiation was started at approximately 90% confluency of hPSCs. The basal media (termed N2B27) used for the differentiation consisted of DMEM/F12 medium (Gibco, Cat. 31331-028) and Neurobasal (Gibco, Cat. 21103-049) in a 1:1 ratio, supplemented with 0.5x N2 (Gibco, Cat. 17502-048), 0.5x B27 (Gibco, Cat. 17504-044), 1x GlutaMAXTM (Gibco, Cat. 35050038), and 1x Penicillin-Streptomycin (Sigma-Aldrich, Cat. P0781-100ML).

For the FGF8 strategy we modified our previously published GnRH neuron differentiation protocol[18]. First, cells were subjected to dual SMAD inhibition (dSMADi) for 10 days by adding 10 µM SB431542 (Sigma, Cat. S4317) and 2 µM dorsomorphin (Selleckchem, Cat. S7306) to N2B27 media, which directs hPSCs to a neural stem cell fate[18]. Cells were then split at a 1:2 ratio on day 10 using collagenase IV (Thermo Fisher, Cat. 17104-019) for 13 min and kept in N2B27 media supplemented with 10 µM Y-27632 (Selleckchem, Cat. S1049) overnight. Next, cells were maintained in N2B27 media supplemented with 100 ng/mL of FGF8b (PeproTech, Cat. AF-100-25-250UG) for 30 days, producing anterior neural progenitor cells (NPs). The cells were split twice: first, on day 20 at 1:8 ratio using EDTA for 8 min and plating them on N2B27 with FGF8b; and second, on day 40 at 1:2 ratio, using EDTA for 4 min and plating them on N2B27 with FGF8b. Finally, cells were switched to Brainphys media (Stemcell Technologies, Cat. 05790) supplemented with 10 µM DAPT (Selleckchem, Cat. S2215) to induce Notch inhibition and neuron emergence, and with 10 ng/mL BDNF (Bio-Techne, 11166-BD) to promote neuron maturation. Cell culture media was fully refreshed every day until day 40, after which only half-media changes were performed. Neurons were collected on day 45.

For the SHH strategy, we modified a previously published hypothalamic differentiation protocol[19]. hPSCs were differentiated in N2B27 media. They were subjected to dSMADi for 8 days using 100 nM LDN-193189 (Selleckchem, Cat. S2618) and 10 µM SB431542. Concomitant SHH-induced ventralization was perfomed via addition of the SHH agonists smoothened agonist (SAG; MilliporeSigma, Cat. 56-666-01) and purmorphamine (PM; MedChemExpress, Cat. HY-15108) at 1 µM between days 2 to 8. On day 9, the media was switched to N2B27 supplemented with 10 µM DAPT for 7 days. Cells were split on day 12 using TrypLE™ Express (Gibco, Cat. 12-605-010) for 4 min, centrifuging for 4 min at 250 rcf, and resuspending in N2B27 media supplemented with 10 µM Y-27632. Cells were plated on matrigel-coated plates at a density of 1*10^6^ cells/dish; after 4 h, the media was refreshed to N2B27 supplemented with DAPT. On day 16, neuron maturation was induced by applying 20 ng/mL of BDNF. Cells were collected on day 45. The culture media was refreshed daily until day 17, after which half media changes were performed every other day.

### Real time quantitative PCR (RT-qPCR)

Following the manufacturer’s instructions, RNA isolation was carried out using the NucleoSpin RNA plus kit (Machery-Nagel, Cat. 740984). 1 µg of RNA was reverse-transcribed into cDNA using iScript™ cDNA synthesis kit (BIO-RAD, Cat. 1708890). 1x HOT FIREPol® EvaGreen® qPCR Mix Plus (no ROX) (Solis BioDyne, Cat. 08-25-00001) was used to perform RT-qPCR, which was measured with a CFX Opus 96 Dx machine (Bio-Rad). Relative gene expression was calculated using the 2^-ΔΔCt^ method by normalizing to the expression of the housekeeping gene *PPIG*. Statistical analyses were performed with -ΔΔCt values to avoid inherent variation of non-log scale values[20]. Supplementary Table 1 contains the primers utilized in this study.

## Results

We followed two distinct strategies to differentiate neuron cultures expressing *KISS1*, according to the postulated different neurodevelopmental origins of the Kiss^ARC^ and Kiss^POA^ populations[14]. First, we hypothesized that anterior neural progenitors that give rise to GnRH neurons may receive similar developmental signals to the progenitors of Kiss^POA^ neurons, given their shared expression of markers such as *FOXG1* and *FGFR1*[14,15,17]. Following this logic, we first set to develop a differentiation strategy that would induce the expression of such factors in the cells[21,22]. Hence, we differentiated hPSCs according to our previously-published GnRH neuron protocol encompassing dSMADi, FGF8b treatment, and Notch inhibition[18]. The duration of the FGF8b treatment was modified by extending it for 30 days to enable the emergence and proliferation of more neural progenitors, and in the final phase of the protocol, a neuronal maturation medium was used (Fig. 1a). This strategy resulted in a significant increase in *KISS1* expression at the end of the protocol (day 45), compared to undifferentiated hPSCs (Fig. 1b). We termed this strategy “FGF8 protocol”.

**Fig. 1.**
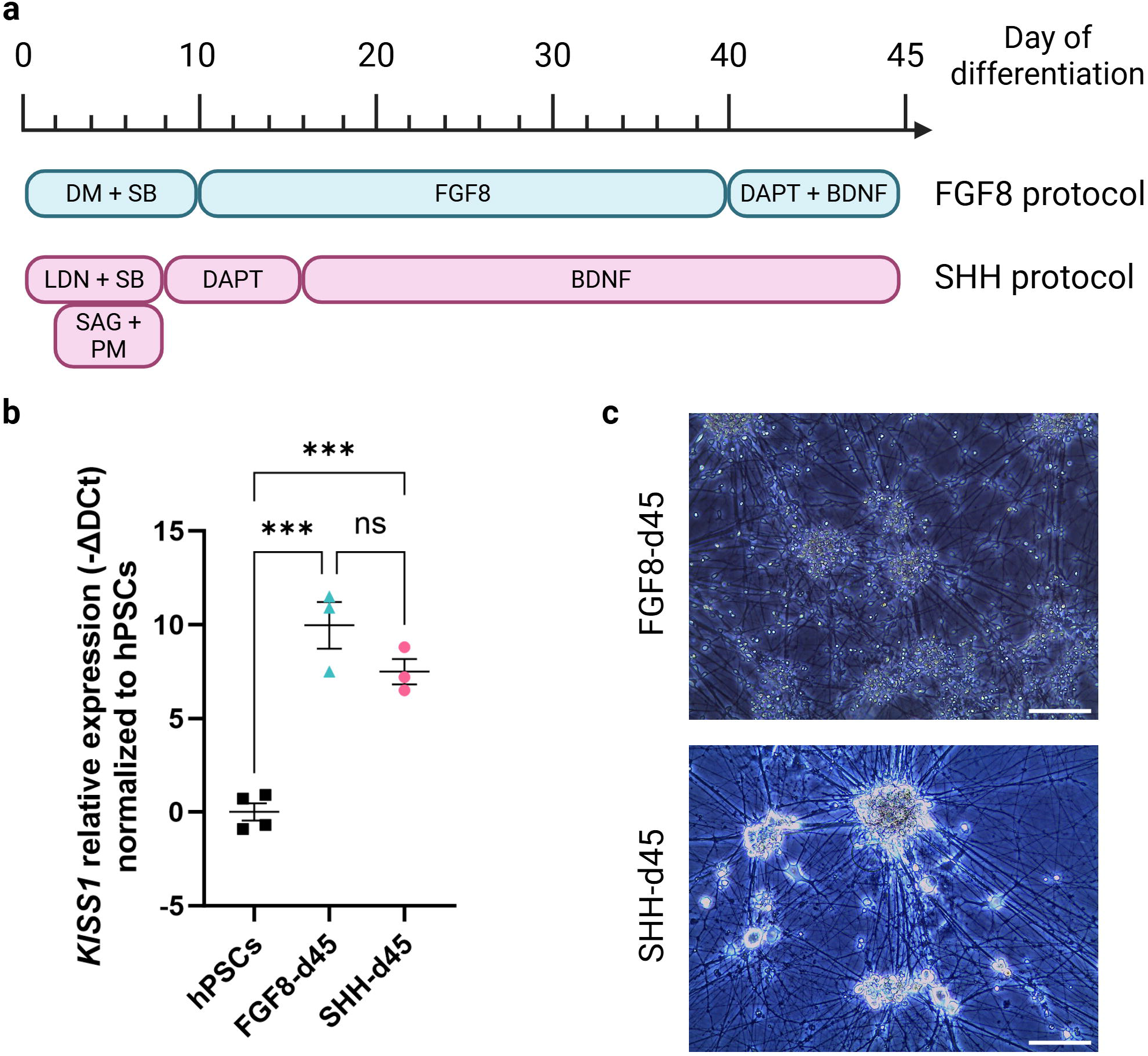
Kiss neuron differentiation strategies. a) Schematic of the two differentiation strategies, FGF8 protocol and SHH protocol. b) *KISS1* expression on day 0 (n = 4) and day 45 of the protocols (n=3). Graphs represent mean ± SEM. Significance was tested with one-way ANOVA. c) Neuron morphology at day 45 of the differentiation protocols. Scale bars represent 100 μm.

Next, following the logic that Kiss^ARC^ neurons can emerge from *POMC*-expressing progenitors[9,10,23], we modified an existing hypothalamic differentiation protocol that directs hPSCs to POMC neurons consisting of dSMADi, SHH activation, and Notch inhibition[19] (Fig. 1a). We observed significant *KISS1* expression in the neuron cultures at day 45 compared to undifferentiated hPSCs (Figure 1b). We termed this strategy “SHH protocol”.

*KISS1* expression was not significantly different between our two differentiation strategies (Fig. 1b). Average raw Ct values (before normalization) for all biological repeats were 35.8 for hPSCs, 26.8 for FGF8-d45, and 29.0 for SHH-d45. Neuron morphology at day 45 from both protocols can be observed in Figure 1c.

We then assessed the expression of *NKX2-1*, a key ventralization factor, at two distinct time points of each differentiation protocol: in neural progenitors corresponding to day 12 of the SHH protocol (SHH-d12) or day 20 of the FGF8 protocol (FGF8-d20); and in differentiated neurons on day 45 of the SHH (SHH-d45) and FGF8 (FGF8-d45) protocols. SHH-d12 and SHH-d45 both showed significant induction of *NKX2-1* compared to undifferentiated hPSCs (Fig. 2a). The high expression of *NKX2*-1 was sustained in day 45 cultures, not differing from SHH-d12 significantly (Fig. 2a). On the contrary, FGF8-d20 and FGF8-d45 cells showed significant but low *NKX2*-1 induction, with *NKX2*-1 expression increasing in FGF8-d45 neurons compared to FGF8-d20 progenitors (Fig. 2b). While SHH-d12 showed significantly higher *NKX2-1* expression than FGF8-d20, SHH-d45 and FGF8-d45 did not show a significant difference in expression. This suggested different temporal expression dynamics of *NKX2*-1 in the protocols.

**Fig. 2.**
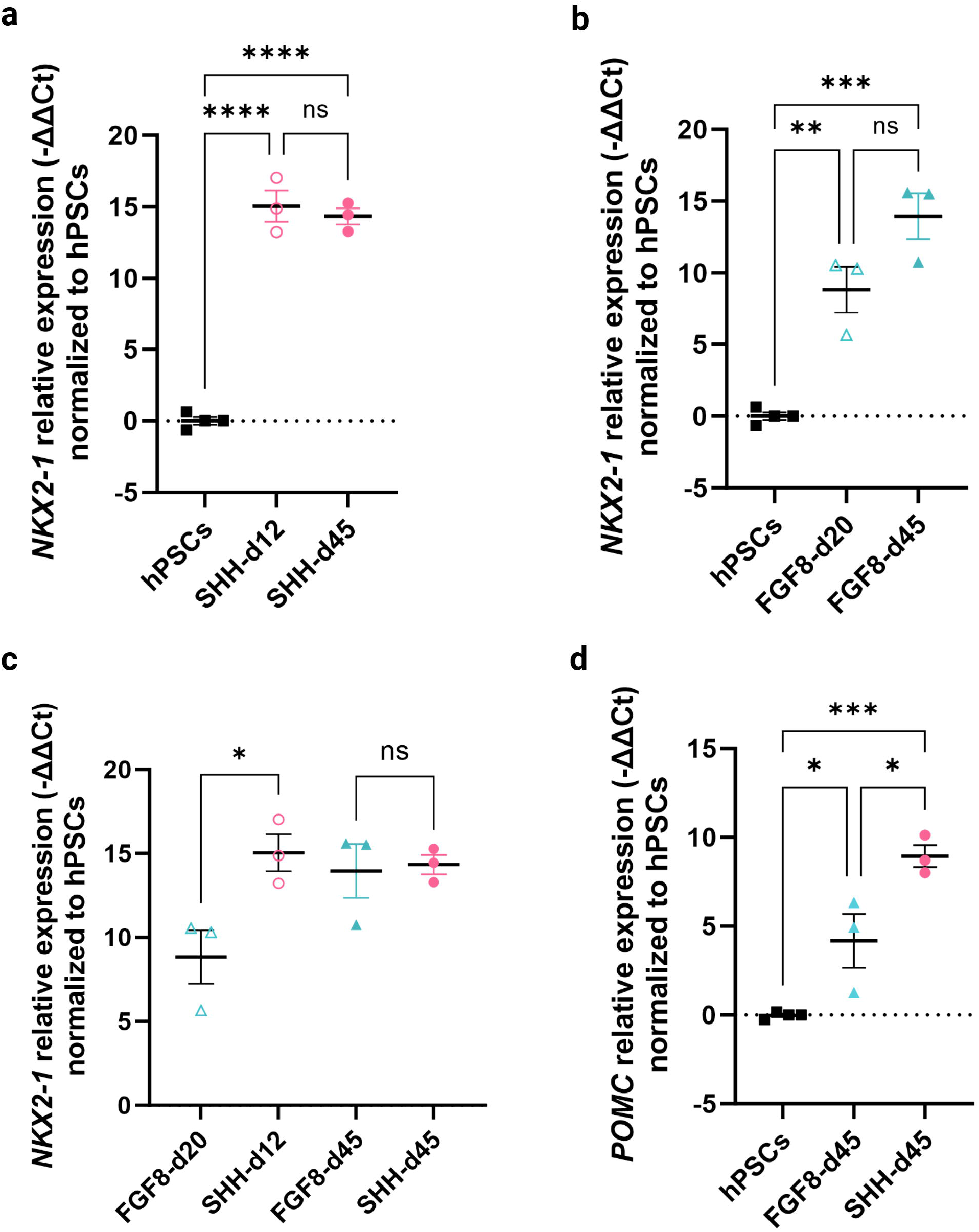
*NKX2*-1 and *POMC* expression in the Kiss differentiation protocols. a) *NKX2-1* expression during the SHH protocol (n = 4 for hPSCs, n = 3 for day 12, n = 3 for day 45). b) *NKX2-1* expression during the FGF8 protocol (n = 4 for hPSCs, n = 3 for day 20, n = 3 for day 45). c) *NKX2*-1 expression comparison between neural progenitors of the SHH protocol (SHH-d12) and FGF8 protocol (FGF8-d20), and between neuron cultures on day 45 of the SHH and FGF8 protocols (d45) (n = 3 for all). d) *POMC* expression on day 45 neurons generated with the FGF8 and SHH protocols (n = 4 for hPSCs, n = 3 for day 45). Graphs represent mean ± SEM. Statistical significance was tested with one-way ANOVA for all graphs.

We additionally assessed the expression of *POMC* in the neural cultures and confirmed that neuron cultures differentiated under the SHH protocol exhibited significant *POMC* expression compared to undifferentiated hPSCs (Fig. 2d). *POMC* expression in the FGF8 protocol was low and significantly diminished compared to neurons derived with the SHH protocol (Fig. 2d).

Finally, other markers previously associated with Kiss^ARC^ and/or Kiss^POA^ neurons were checked in the day 45 neuron cultures with RT-qPCR (Fig. 3). Compared to the FGF8 protocol, SHH-derived cultures displayed higher expression of the ARC markers *NHLH2* and *NR5A2*, both of which have been implicated in Kiss^ARC^ neuron development and function in mice[12,24,25]. Meanwhile, FGF8 protocol-derived cultures showed significantly higher expression of *FOXG1*, an anterior telencephalon marker shown to be differentially expressed in Kiss^POA^ neurons compared to Kiss^ARC^ neurons[14]. However, *SOX14*, which has been implicated in Kiss^ARC^ development[25], was similarly expressed in cultures differentiated with both strategies. Expression of *PENK* and *FGFR1* did not differ signifiantly between cultures differentiated with the SHH and FGF8 protocols. *TAC3*, the gene encoding NKB in humans that is part of the KNDy identity[4], showed a trend of higher expression in the SHH protocol, though this result was not significant.

**Fig. 3.**
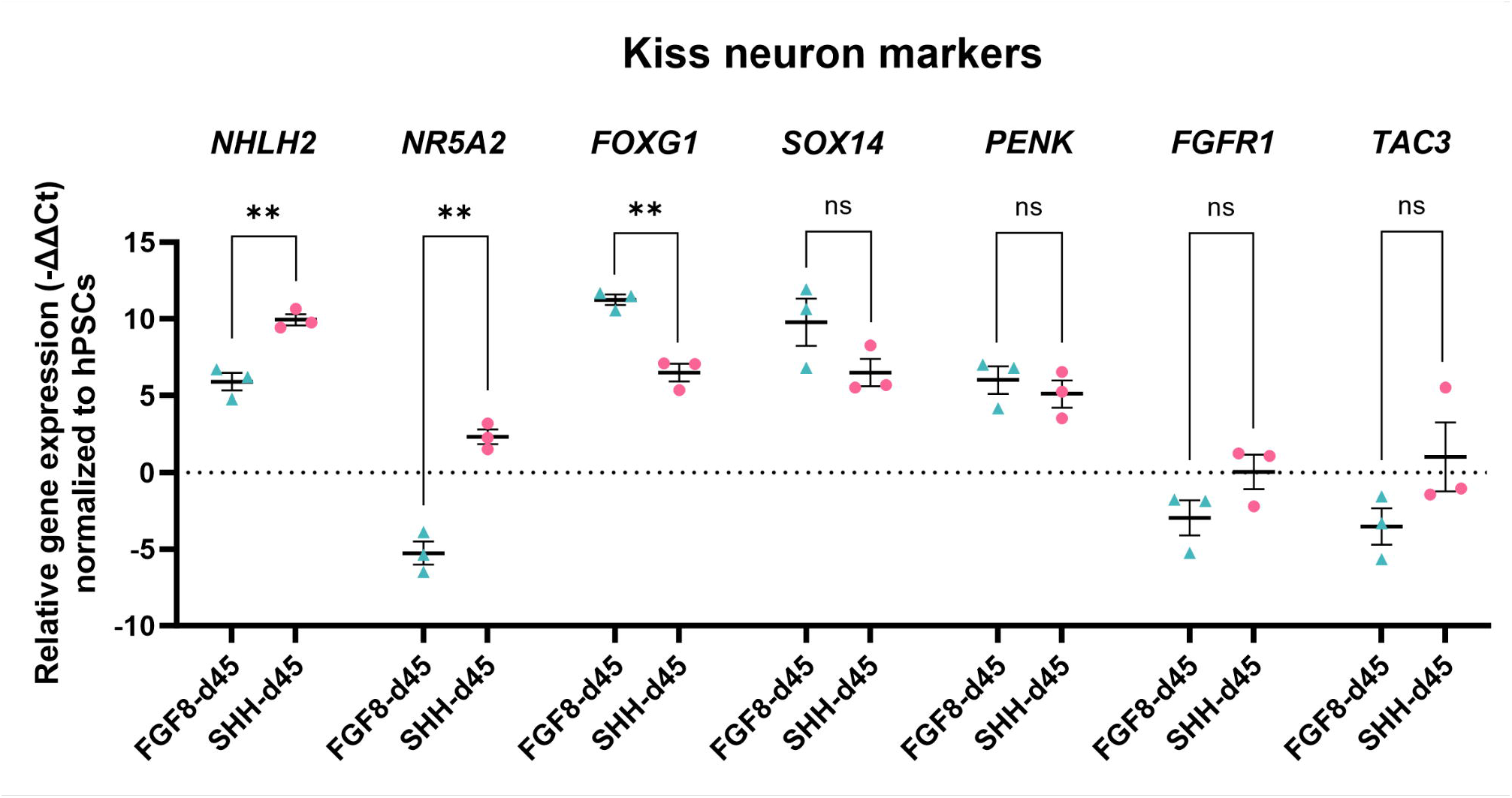
Expression of different Kiss neuron markers in the FGF8 and SHH differentiation protocols on day 45 (n = 3 for all). Graphs represent mean ± SEM. Statistical significance was tested with one-way ANOVA.

## Discussion

Here, we provide with the first study to show successful differentiation of hPSC-derived neuron cultures expressing *KISS1* using a monolayer cell culture strategy. These results show that *KISS1* expression can be achieved with a combination of TGFb/SMAD2/3 inhibition (dSMADi), ventralization via SHH activation, and Notch inhibition. This is in agreement with a recent publication that differentiated Kiss neurons from mouse embryonic stem cells employing SHH activation and Notch inhibition[26]. Indeed, SHH-related ventralization induces *NKX2-1*[27], which enhances hypothalamic specification and ARC patterning[28–30] and which is required for Kiss neuron function[31]. As such, we found *NKX2-1* was highly expressed in our SHH-derived neuron cultures. These cultures also expressed *NHLH2* and *NR5A2*, which are markers of ARC neurons including Kiss and POMC neurons[12,24,32].

On the other hand, TGFb/SMAD2/3 inhibition followed by prolonged FGF8b exposure resulted in significant *KISS1* expression. This may stem from the anteriorization promoted by these factors, which induce the expression of *FOXG1* and *DLX1/2* in neural progenitors and neurons[21], genes that have been found to be differentially expressed in mouse Kiss^AVPV^ compared to Kiss^ARC^ neurons[14]. Indeed, our FGF8-derived neuron cultures induced high *FOXG1* expression. Moreover, *FGFR1* expression is differentially upregulated in mouse Kiss^AVPV^ neurons[14], suggesting involvement of FGF8-FGFR1 signaling in Kiss neuron development. In our work, *KISS1* expression was observed in the FGF8 protocol in the absence of SHH activation, suggesting that ventralization via SHH signaling is not necessarily required for Kiss^POA^ neuron development. Interestingly, *NKX2-1* was expressed in these neuron cultures on day 45. This is consistent with previous literature showing that murine Kiss^AVPV^ neurons coexpress NKX2-1[33]. *FGF8* is expressed in the rostral prosencephalon in chicken at HH12, and is followed by the appearance of *NKX2-1* in the rostroventral telencephalon, which later acquires several *SHH* domains that interact with *FGF8* domains[34]. This data, together with our findings, suggest that FGF8 may act as an early patterning factor in the AVPV/POA, leading to permissive *NKX2-1* expression and eventually Kiss^POA^ neuron emergence.

Interestingly, we found no significant difference in the expression of several markers, including *SOX14, PENK*, and *FGFR1*. While PENK has been associated mainly with Kiss^POA^ neurons, it has also been found expressed in human Kiss^ARC^ neurons[7]. Similarly, *FGFR1* has also been found expressed in mouse Kiss^ARC^ neurons, with its expression changing with estrogen feedback[35].

It is tempting to hypothesize that our two approaches produce Kiss neurons with different properties (ARC vs POA), but further characterization is required to validate this hypothesis. Currently, the most important limitation is that neuron cultures likely contain more than one type of neurons (e.g. POMC neurons in the Kiss^ARC^ cultures) and thus RT-qPCR assays are interpreted as the average gene expression of the whole neural population, not just of Kiss-expressing cells. Further studies using immunocytochemistry and single-cell RNA-sequencing will help characterization of hPSC-derived Kiss neurons in depth.

In conclusion, we present the first evidence that neuron cultures that express significant *KISS1* can be differentiated from hPSCs. These results pave the way to further characterization of Kiss neuron types and provide a tool to explore the heterogeinity in the human Kiss neural system.

## Statements

### Statement of Ethics

Established human cell line H9 embryonic stem cells used in this study are registered in the EU, under the license of WiCell®. No animal or patient experiments were conducted in this research.

## Acknowledgement

A preprint version of this article is available on BioRxiv.

## Conflict of Interest Statement

The authors have no conflicts of interest to declare.

## Funding Sources

Open Access funding was provided by University of Helsinki (including Helsinki University Central Hospital). This study was financially supported by the Research Council of Finland (368584), Sigrid Juselius Foundation, Foundation for Pediatric Research, The Hospital District of Helsinki and Uusimaa/Children and Adolescents, Vilho, Yrjö ja Kalle Väisälän rahasto. The funders had no role in the design, data collection, data analysis, and reporting of this study.

## Author Contributions

Celia Gomez-Sanchez, MSc: conceptualization, experiments (cell differentiation, RT-qPCR), writing (original draft and review and editing), funding acquisition. Nazli Eskici, PhD: conceptualization, experiments (cell differentiation), writing (review and editing). Shrinidhi Madhusudan, MSc: experiments (cell differentiation, RT-qPCR), writing (review and editing). Karoliina Tanner, BSc: experiments (RT-qPCR). Kirsi Vaaralahti, PhD: writing (review and editing), project administration, funding acquisition. Kristiina Pulli, PhD: writing (review and editing), supervision. Taneli Raivio, MD: conceptualization, writing (review and editing), project administration, supervision, funding acquisition. All authors read and approved of the final manuscript.

## Data Availability Statement

All data generated or analyzed during this study are included in this article. Further enquiries can be directed to the corresponding author.

## Notes

### Competing Interest Statement

The authors have declared no competing interest.

