## Supplementary Table 1 for "Differentiation of KISS1-Expressing Cells from Human Pluripotent Stem Cells: Many Roads To Rome"

**Supplementary Table 1****: RT-qPCR primers used in this study.**

| **Target gene** | **Forward sequence** | **Reverse sequence** |
| --- | --- | --- |
| *FGFR1* | GGAAGGACTCCACTTCCACA | GTCACAGCCACACTCTGCAC |
| *FOXG1* | CCGCACCCGTCAATGACTT | CCGTCGTAAAACTTGGCAAAG |
| *KISS1* | CAAGCCTCAAGGCACTTCTA | AAAGTGGGTGGCACAGAG |
| *NHLH2* | GTCCGGACTCAGCATCATTT | GGAATCTCCCCTCGCTATTC |
| *NKX2-1* | AACCAAGCGCATCCAATCTCAAGG | TGTGCCCAGAGTGAAGTTTGGTCT |
| *NR5A2* | CTTTGTCCCGTGTGTGGAGAT | GTCGGCCCTTACAGCTTCTA |
| *PENK* | GCTTGCGTAATGGAATGTGAAG | TCTTGAGGAAGCTCTGGTTTG |
| *POMC* | CAGGCACTTGCTGGATTCTC | GTTGCTTTCCGTGGTGAGGT |
| *PPIG* | ACTCCCAGCCTGCTTCATAC | TACGTCTGAAACGATCCCTTG |
| *SOX14* | TACGTGGTGCCCTGTAACTG | GGGTCTATGCCAGTCTTGGT |
| *TAC3* | GGATCATGCTGCTATTCACAG | ATGCATGTCACGTTTCTCGG |
